# Supervised learning with word embeddings derived from PubMed captures latent knowledge about protein kinases and cancer

**DOI:** 10.1101/2021.06.11.447943

**Authors:** Vida Ravanmehr, Hannah Blau, Luca Cappelletti, Tommaso Fontana, Leigh Carmody, Ben Coleman, Joshy George, Justin Reese, Marcin Joachimiak, Giovanni Bocci, Carol Bult, Jens Rueter, Elena Casiraghi, Giorgio Valentini, Christopher Mungall, Tudor Oprea, Peter N. Robinson

## Abstract

Inhibiting protein kinases (PKs) that cause cancers has been an important topic in cancer therapy for years. So far, almost 8% of more than 530 PKs have been targeted by FDA-approved medications and around 150 protein kinase inhibitors (PKIs) have been tested in clinical trials. We present an approach based on natural language processing and machine learning to the relations between PKs and cancers, predicting PKs whose inhibition would be efficacious to treat a certain cancer. Our approach represents PKs and cancers as semantically meaningful 100-dimensional vectors based on co-occurrence patterns in PubMed abstracts. We use information about phase I-IV trials in ClinicalTrials.gov to construct a training set for random forest classification. In historical data, associations between PKs and specific cancers could be predicted years in advance with good accuracy. Our model may be a tool to predict the relevance of inhibiting PKs with specific cancers.

## INTRODUCTION

Protein phosphorylation is one of the most important post translational modifications. The human genome encodes 538 protein kinases (PKs), many of which are associated with cancer initiation or progression. Kinases transfer a γ-phosphate group from ATP to serine, threonine, or tyrosine residues; the genome encodes roughly 200 phosphatases that remove a phosphate group from a protein. Protein phosphorylation and dephosphorylation represent an important regulatory mechanism that is involved in virtually every basic cellular process including proliferation, cell cycle, apoptosis, motility, growth, and differentiation. Many PKs promote cell proliferation, survival and migration, and misregulation of kinase activity is a common feature of oncogenesis (1–3). Molecularly targeted cancer therapies are rapidly growing in importance for the treatment of many types of cancer. Many targeted therapies, including small-molecule kinase inhibitors and monoclonal antibodies, act as protein kinase inhibitors (PKIs). Since the introduction of the initial PKI in the 1980s, at least 37 PKIs have received FDA approval for cancer therapy and over 150 kinase-targeted drugs are in clinical trials (3).

PKIs are not equally effective for all cancer types; instead, specific characteristics of each tumor, including genetics, tumor microenvironment, drug resistance, and pharmacogenomics determine how useful a compound will be in the treatment of a given cancer. Factors including whether a particular kinase exhibits activating mutations in a given cancer, or whether downstream targets of the kinase are mutated strongly influence the susceptibility of a cancer to a given PKI. Characteristics of pathways related to those mutated in a given cancer can also influence response to targeted treatment (4). In addition, most PKIs target more than one protein with a range from highly to poorly selective (5). It is therefore not always possible to reliably predict whether a given PKI will be efficacious against a given type of cancer. For instance, imatinib, which targets BCR-ABL, c-ABl, PDGFR, and c-Kit, was found not be effective in uveal melanoma despite high expression of KIT, an unexpected finding that was interpreted to be related to the lack of ERK phosphorylation in these tumors (6).

In this work, we pose the question of whether one can use knowledge latent in the published literature to predict whether inhibition of a given PK is an effective treatment of a cancer. Correct predictions could be used to prioritize clinical trials of a cancer with PKIs that target the PK in question. We present an algorithm that embeds cancer-relevant concepts in PubMed abstracts into a semantically meaningful vector space. Difference vectors between all pairs of cancers and PKs are formed by subtracting each cancer vector from each PK vector. Only some of the resulting vectors are expected to represent valid analogies in the sense that inhibition of the kinase is helpful in treating the cancer. Data from ClinicalTrials.gov are used to form a training set for random forest classification.

In this work, we explore machine learning approaches toward the prediction of the clinical relevance of PKs for specific cancers so that inhibiting the PKs may lead to treating the cancer. In particular, our aim is to exploit the large corpus of clinical text data available in PubMed abstracts to discover novel associations between PK and cancer, leveraging natural language processing approaches based on word embedding that have been successfully applied to text analysis, representation and classification tasks (7). On historical data, we achieved an area on the receiver operating characteristics (ROC) curve (AUROC) of up to 90% for predicting successful trials of all phases and we achieved AUROC up to 95% for predicting successful phase IV trials. Predictions based on PubMed data up to 2020 revealed 1432 PK-cancer pairs with above-threshold probabilities.

## RESULTS

We developed a machine learning approach towards leveraging knowledge latent in the published literature to predict pairs of PKs and cancers that will be the subject of clinical trials published in the ClinicalTrials.gov resource. Our pipeline assigns embeddings to words and concepts in the original texts, extracts embeddings related to cancers and PKs, and applies random forest classification to predict pairs of cancer and PKs that correspond to clinical trials in which a PKI that inhibits the PKs is used to treat a given form of cancer.

To this end, we extracted PubMed articles from 1939 to 2020 (with a gap of seven years from 1940 to 1946) based on their MeSH descriptions for neoplasms and PKs, obtaining 2,779,507 relevant articles on the basis of 698 MeSH terms for neoplasms and 218 MeSH terms for PKs. We first prepared the abstract texts for word embedding by concept replacement, stop word removal, and lemmatization (**Figure 1A**). The preprocessing step has several desirable effects. Firstly, it merges synonyms; for instance, “breast cancer” and “Cancer of Breast” are both replaced by the corresponding concept id, MESHD001943. Lemmatization replaces inflected word forms with a common base form, for instance “higher” is replaced by “high” in the example of **Figure 1A**. Stop words, i.e., common words such as ‘a’ and ‘and’, are removed because they do not carry much semantic information. All punctuation marks such as “,” and “.” are removed and all letters are converted to lowercase.

**Figure 1.**
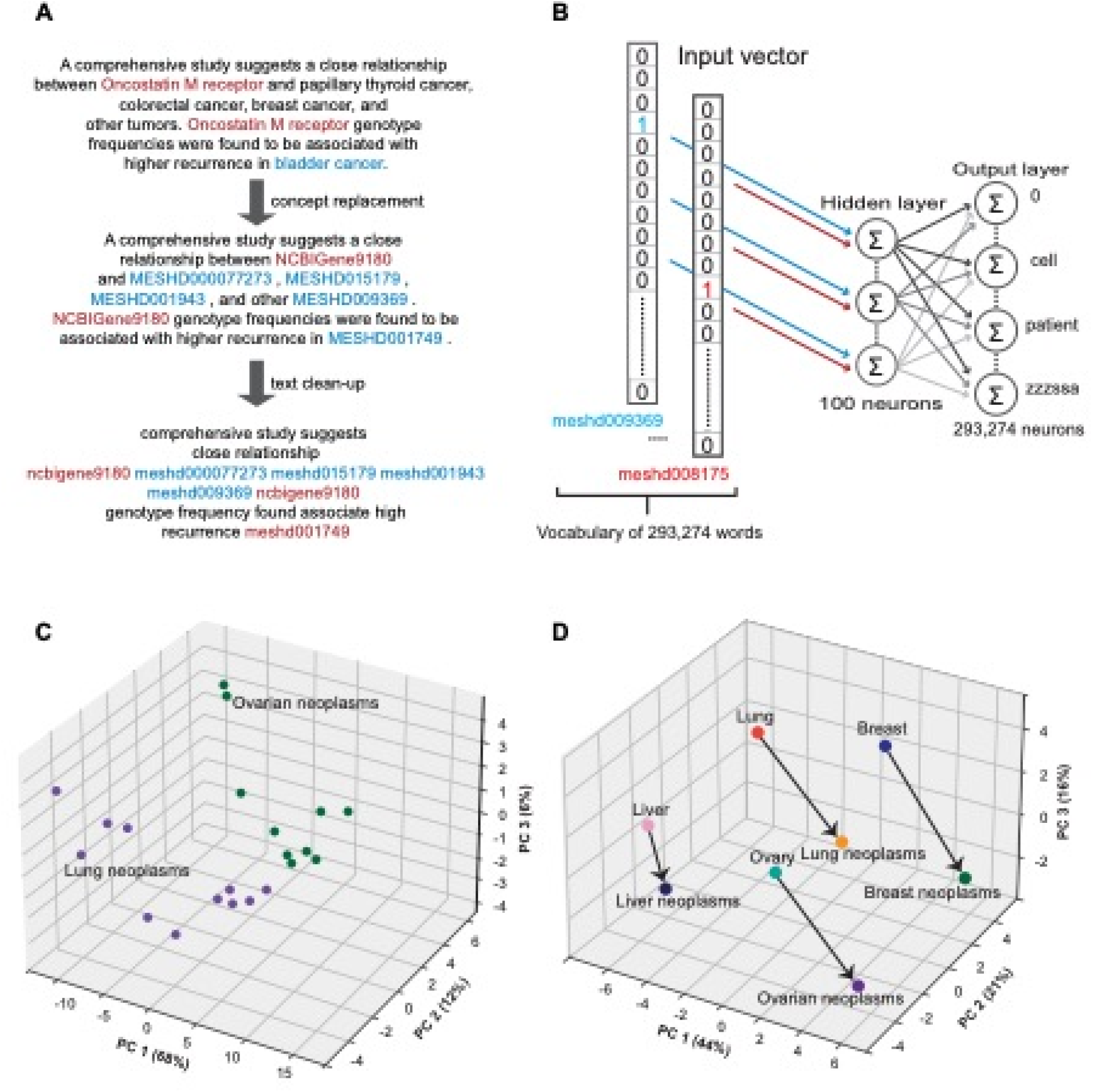
Overview of concept embedding algorithms. **A)** An example of preprocessing on a text. **B)** Word2vec skip-gram learning. Words (potentially replaced by concept IDs) are transformed from a one-hot representation into a low-dimensional vector through a one hidden-layer neural network trained to predict context words. Backpropagation learning adjusts the weights of the hidden layer whose output can be interpreted as a low dimensional semantic representation (word vector) of the one-hot encoded input word. The output layer contains probabilities of a word to occur at a neighboring position to the target word. **C)** Word vectors are represented in the space induced by the first three principle components. Vectors representing Lung Neoplasms and descendent terms are shown in purple; vectors representing Ovarian Neoplasms and descendent terms are shown in green. **D)** Positions of 8 vectors in three-dimensional PCA space are shown. Arrows are used to connect pairs of vectors representing tissues and cancer that affects the tissue. It can be seen that the pairs form analogies such that, for instance, *f* (*Lung*) - *f* (*Lung neoplasms*) ≈ *f* (*Breast*) - *f* (*Breast neoplasms*), where *f(*.) represents the embedding of a word in the vector space.

Following this, word embedding was performed with a skip-gram model (**Figure 1B**). This step creates 100-dimensional vector representations (embeddings) of the words and concepts of the processed abstract texts. The motivating idea of the word2vec algorithm is that because words with similar meanings often appear together, the corresponding embeddings will be located close to each other in the vector space (8). In addition, word vectors may reflect semantic relationships between words in ways that can be expressed as analogies, e.g., France is to Paris as Germany is to Berlin (9). In our data, embeddings for ovarian neoplasms and lung neoplasms formed two distinct clusters (**Figure 1C**). Additionally, we identified pairs of vectors that demonstrated the semantic relation “organ is to organ-specific cancer” (**Figure 1D**).

### PKIs targeting PKs

The goal of our approach is to predict clinical studies related to therapeutically relevant PK-cancer pairs. To do so, we curated information available in DrugCentral (10), and identified 75 PKIs that have been used to treat cancers. In many cases, the PKIs inhibit multiple PKs at a < 0.3μM cutoff, and a total of 84 PKs are inhibited by these kinases. The mean number of PKs inhibited by a given PKI was 2.8 (median 2, min. 1, max. 5), and the mean number of PKIs that inhibit a given PK was 2.5 (median 2, min. 1, max. 13) (**Supplementary Material Figures S1A and S1B**).

We retrieved clinical studies that involved these PKIs from the ClinicalTrials.gov resource (11), identifying 2105 phase I, 3185 phase II, 555 phase III, and 217 phase IV studies performed between 1991 and 2021 (Total 6062; **Supplementary Material Figures S2A and S2B**).

### Random forest classification of PK-cancer pairs

We then used the word embeddings as the basis for machine learning classification. We first extracted the 698 embeddings representing neoplasms and the 218 embeddings for PKs. For a list of the 75 PKIs that have been used to treat cancers, we extracted information from DrugCentral regarding the PKs that are inhibited by each PKI with the highest affinities (see Methods for details). We then extracted data from ClinicalTrials.gov about clinical trials in which the use of the PKI to treat a certain cancer was investigated. We interpret a phase IV (postmarketing) trial as evidence that the PKI demonstrated efficacy in treating the cancer. **Figure 2B** offers an example of how our procedure would associate EGFR with three cancers against which the PKI afatinib demonstrated efficacy.

**Figure 2.**
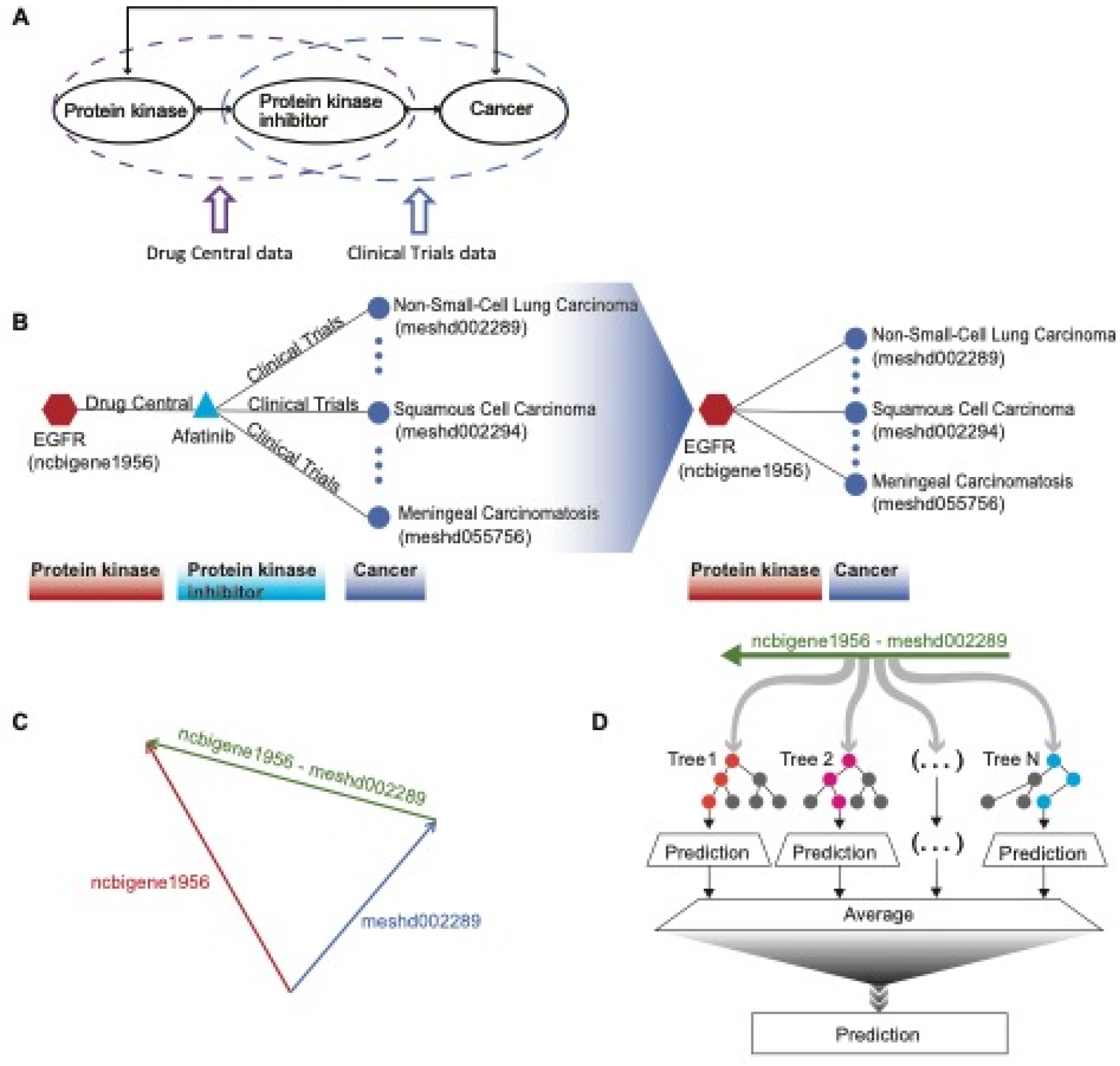
Machine learning to predict PKs relevant for treating cancer. **A)** Information about phase I-IV trials of PKI was derived from ClinicalTrials.gov and DrugCentral. **B)** An example showing how the PK-cancer pairs were derived from ClinicalTrials.gov and DrugCentral. **C)** The embedded vectors derived from skip-gram analysis of PubMed abstracts were used to generate “analogy vectors” by subtracting vectors of cancers from vectors of PKs. Positives were defined by the Clinical Trials data, and negatives were chosen from the remaining vectors. **D)** Random forest classification was performed using analogy vectors as input.

It can be seen from **Figure 1D** that only some pairs of tissues and cancers form valid analogies. For instance, *f* (*Lung*) - *f* (*Lung neoplasms*) ≈ *f* (*Breast*) - *f* (*Breast neoplasms*), while it is not true that *f*(*Lung*) - *f*(*Breast neoplasms*) ≈ *f*(*Breast*) - *f*(*Lung neoplasms*). We reasoned that vectors of the form *f* (*PK*) - *f* (*Cancer*)could be used for classification if the distribution of vectors that are derived from PK whose inhibition can be exploited to treat a given cancer differ from the general distribution of vectors derived from arbitrary pairs of PKs and cancers. For instance, the PKI sorafenib inhibits the kinases RAF, VEGFR 1-3, PDGFR, c-KIT and RET and significantly improves progression-free survival compared with placebo in patients with progressive radioactive iodine-refractory differentiated thyroid cancer (12). For the purposes of our analysis, we define the vectors formed by subtracting the vector for Thyroid Neoplasms (MeSH D013964) and those for RAF, VEGFR 1-3, PDGFR, c-KIT and RET, as belonging to the positive set. We assume that the vast majority of relations between PKs and cancers are not therapeutically relevant in this way, although data that proves this negative role is not generally available in the literature, so we assume that vectors that are not in our positive set are negative.

It is worth noting that several relations between words, including analogy, are approximately preserved by simple linear combinations (e.g. difference) of the vectors representing the words in the embedded space (13). Here, for each PK-cancer pair, we define a difference vector by subtracting the cancer vector from the corresponding PK vector (**Figure 2C**). The sets of positive and negative vectors defined in this way are used for random forest learning, whereby the features used by the random forest are provided by the magnitudes of the vectors in each of the 100 dimensions of the embedded vectors (**Figure 2D**).

As an example of our procedure, we describe the historical validation pipeline for the target year of 2010 in detail. 2533 clinical trials were published in ClinicalTrials.gov between 1991 and 2010, resulting in 132 PK-cancer pairs. Corresponding negative training and test sets were constructed by randomly choosing 1158 PK-cancer pairs not mentioned in the ClinicalTrials.gov data in 2010 or before (see Methods for more details). Random forest classification was trained on the vectors obtained by subtracting vectors corresponding to cancers from vectors corresponding to PKs of the training set. The parameters of the random forest classifier are explained in Methods. In our first analysis, we assessed the performance of classification by evaluating performance based on newly published clinical trials in 2011-2012, 2013-2014, and so on up to 2019-2020. In **Figure 3**, we have provided the ROC curves using PubMed abstracts published up to 2010 (**Figures 3A, 3B**) and PubMed abstracts published up to 2014 (**Figures 3C, 3D**). The number of PK-cancer pairs in positive test sets is shown with *n*. The area under the receiver operating characteristic curve (AUROC) was 77% for data in the first two years immediately following the target year and showed some fluctuations in the next 2-year intervals and reached to 85% in 2015-2016 and 2019-2020 (**Figure 3A**). In our second analysis on historical predictions, we assessed the performance of classification by evaluating performance based on newly published clinical trials in ClinicalTrials.gov in 2011 (1 year after 2010), 2011-2012 (2 years after 2010) and so on up to 2011-2020 (10 years after 2010). The AUROC scores start from 80% in 2011, immediately one year after 2010 and stay within the same range between 78% and 83% over the following time periods, reaching the AUROC score of 82% in the last time period (2011-2020) (**Figure 3B**). We have demonstrated the ROC curves of predictions using PubMed abstracts up to 2012, 2016 and 2018 in **Supplementary Material Figure S3**. In the second part of our experiments, we again considered the same target years, 2010 and 2014. But, instead of predicting PK-cancer pairs corresponding to all phases I, II, III and IV, we only considered prediction of PK-cancer pairs based on phase IV clinical trial studies. The AUROC results are provided in **Figure 4**. For the target year 2014, in some time periods, such as 2015 and 2015-2016 and 2016-2017, there were no PK-cancer pairs based on phase IV clinical trials in the positive test set. So, there are no ROC curves for these time periods. Overall, the prediction of PK-cancer pairs based on phase IV clinical trials shows a gradual decrease in AUROC from the year after the target year to the latest years. For example, predictions based on data in PubMed up to and including 2010 predicted multiple interactions between members of the Neurotrophic Tyrosine Receptor Kinase gene family (NTRK1, NTRK2, NTRK3) and renal cell carcinoma (NTRK1,0.845; NTRK2,0.745; NTRK3, 0.830), hepatocellular carcinoma (NTRK1,0.865; NTRK3, 0.715), breast neoplasms (NTRK1, 0.735; NTRK3, 0.685), Lung Neoplasms (NTRK1, 0.60), and NTRK1 - Gastrointestinal Neoplasms (NTRK1, 0.625) in addition to predictions for Leukemia(NTRK1,0.735, NTRK3,0.725), whereby the prediction probability is shown after the gene. THe three NTRK isoforms are inhibited by entrectinib and larotrectinib. The first mention of the NTRKs, entrectinib and larotrectinib in ClinicalTrials.gov was in 2014. Larotrectinib is a tumor agnostic NTRK3 inhibitor(14) and was approved by the FDA in 2018 (15). Entrectinib induced durable and clinically meaningful responses in patients with NTRK fusion-positive solid tumours(16) and was approved by the FDA in 2019 for solid tumors that have an NTRK gene fusion (17). Additionally, preclinical data showed efficacy for entrectinib in acute myelogenous leukemia (AML) cell lines with endogenous expression of the ETV6-NTRK3 fusion gene (18).

**Figure 3.**
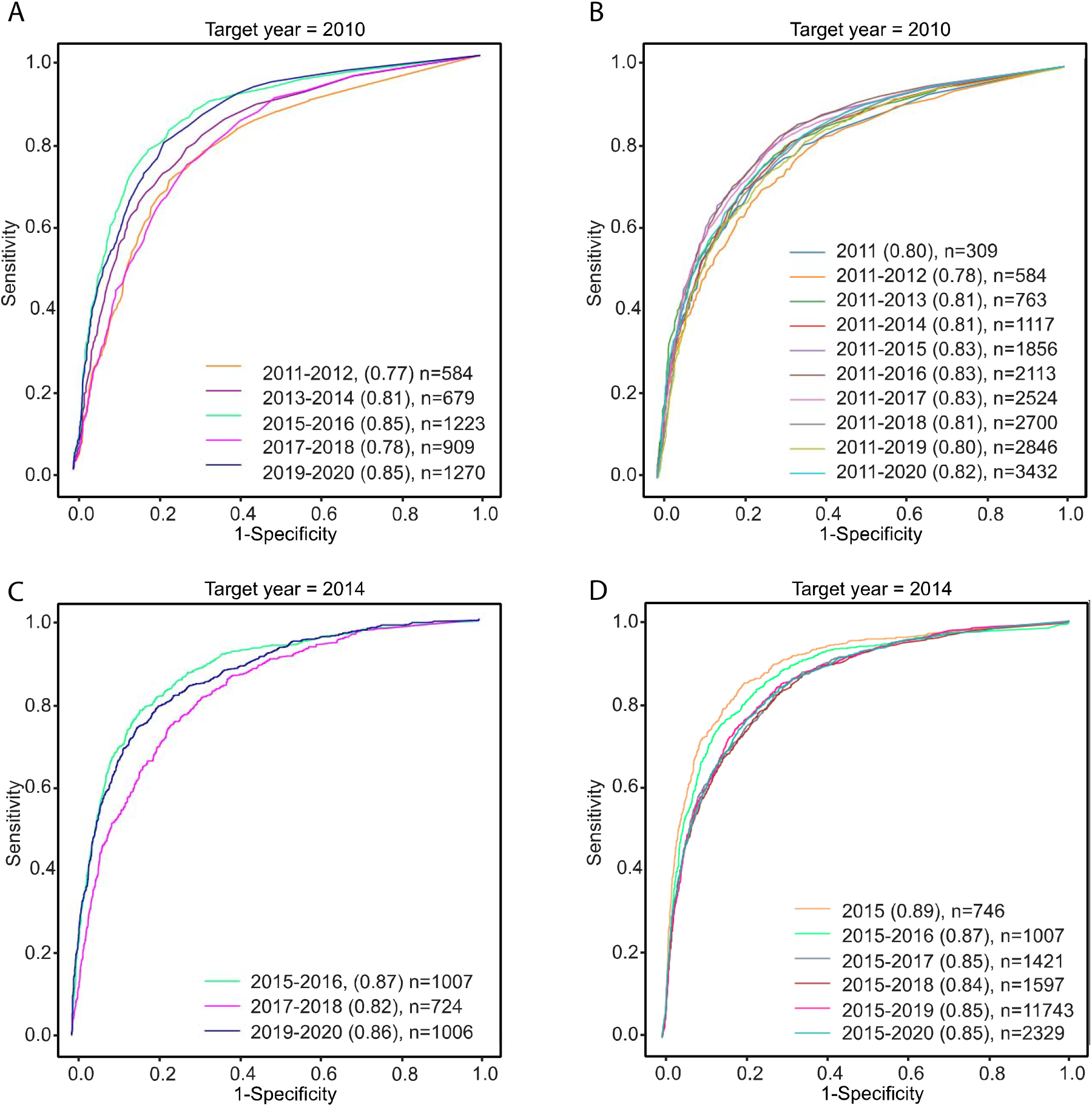
ROC analysis of predicted PK-cancer pairs (all clinical trial phases). **A)** Predictions based on abstracts published up to 2010, grouped according to two-year periods following 2010. **B)** Predictions based on abstracts published up to 2010, grouped according to increasing periods of time in the future. **C)** Predictions based on abstracts published up to 2014, otherwise analogous to panel A. **D)** Predictions based on abstracts published up to 2014, otherwise analogous to panel B.

**Figure 4.**
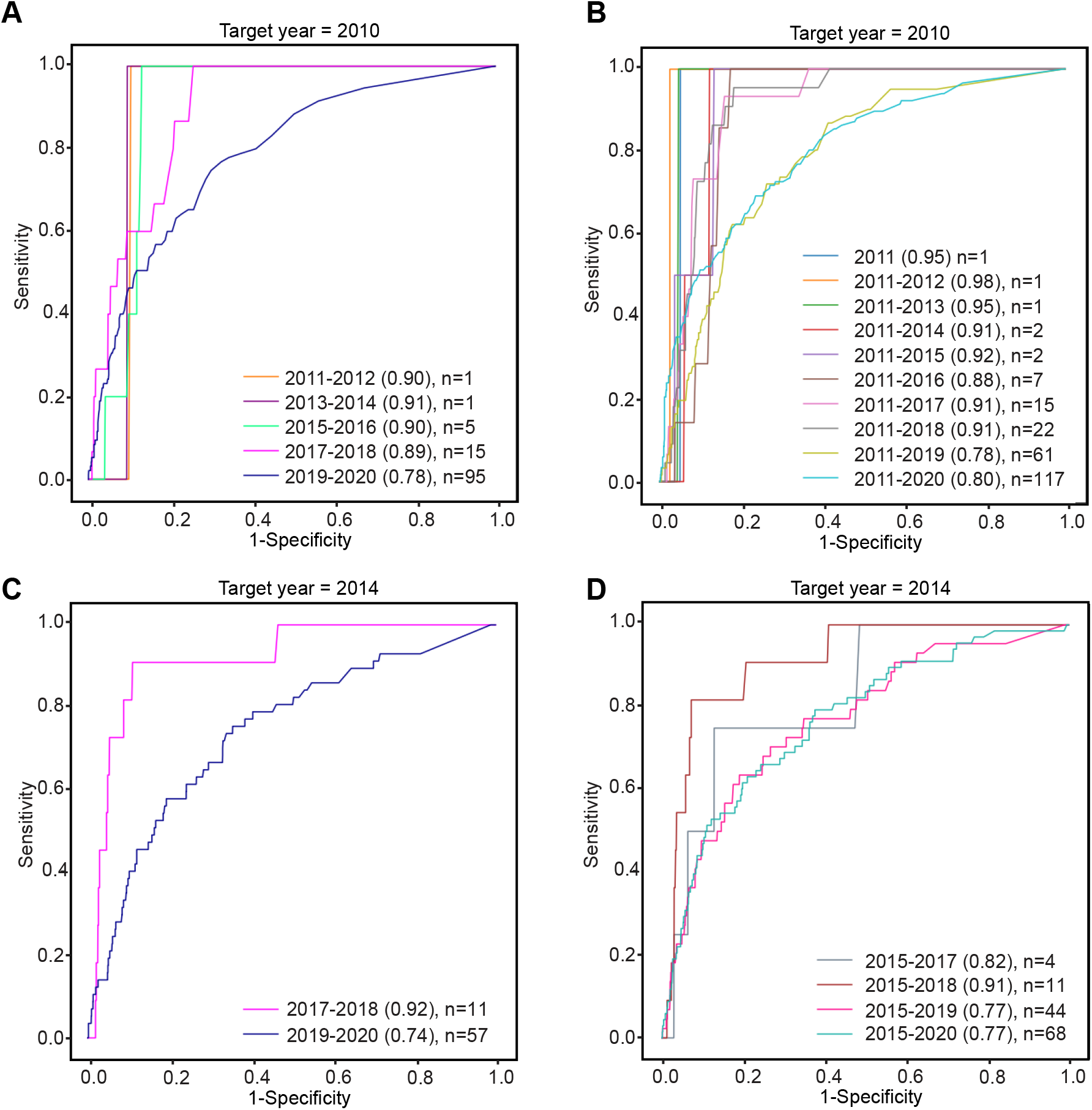
ROC analysis of predicted PK-cancer pairs (phase IV clinical trials). **A)** Predictions based on abstracts published up to 2010, grouped according to two year periods following 2010. **B)** Predictions based on abstracts published up to 2010, grouped according to increasing periods of time in the future. **C)** Predictions based on abstracts published up to 2014, otherwise analogous to panel A. **D)** Predictions based on abstracts published up to 2014, otherwise analogous to panel B.

Finally, we ran our method on our all corpus of PubMed abstracts up to December 2020. We considered all clinical trials up to 2020 and also clinical trials that have been verified in 2021. We then constructed the positive training set using all PK-cancer pairs from clinical trials of phase IV. The negative training set contains randomly generated pairs of PKs and cancers where there was no evidence of treating the cancers by inhibiting the PK in the clinical trials data. Similar to the historical prediction analysis, we chose the size of the negative training set to be 10 times the size of the positive training set. The prediction set includes all possible PK-cancer pairs except those where there was evidence of treating the PKs in any of phase I, II, III or IV clinical trials that have been published so far. The prediction set also contains PK-cancer pairs for PKs that have not been targeted yet. The size of the positive training set, negative training set and prediction set are 559, 5217 and 330922, respectively. The top 10 predictions along with their prediction score from the random forest classifier are given in **Table 1**. Some of these predictions are currently being studied in clinical trials. For those predictions, we provide the phase of the clinical trials along with the ClinicalTrials.gov identifiers of the studies. The comments on predictions for which there is no evidence of any clinical trials are left as blank. Our results in **Table 1** show that the PK *RYK*, which according to our Drug Central data has not been targeted yet, was found to be a potential target in *non-small-cell lung carcinoma*. In **Supplementary Material File 3**, we have provided the predictions with prediction scores at least 0.556. This value was chosen based on the threshold of the AUROC scores at which the true positive rate and false positive rate maximize the g-means, where g-means is the square root of (true positive rate) multiplied by (1-false positive rate).

**Table 1:**
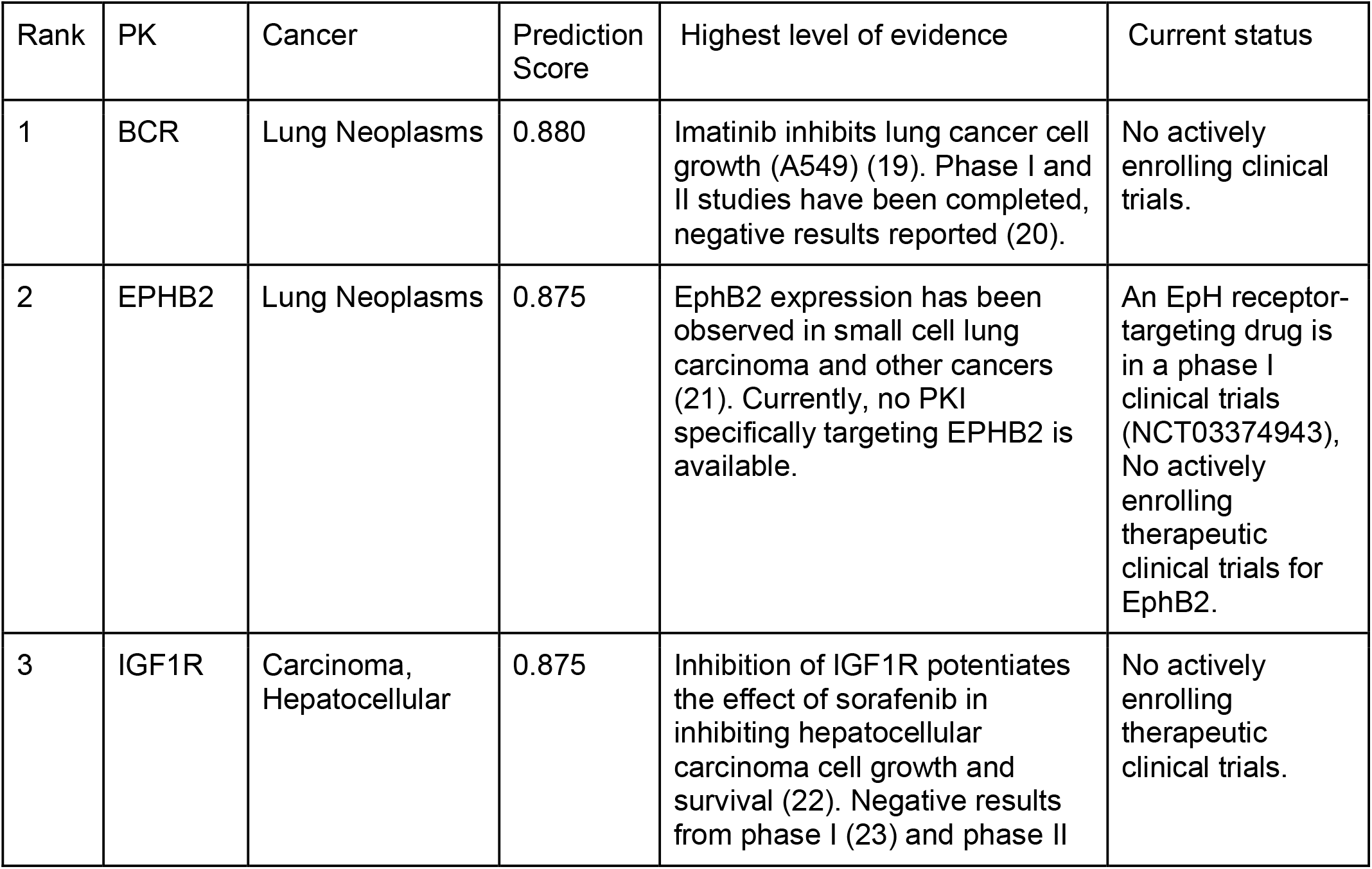
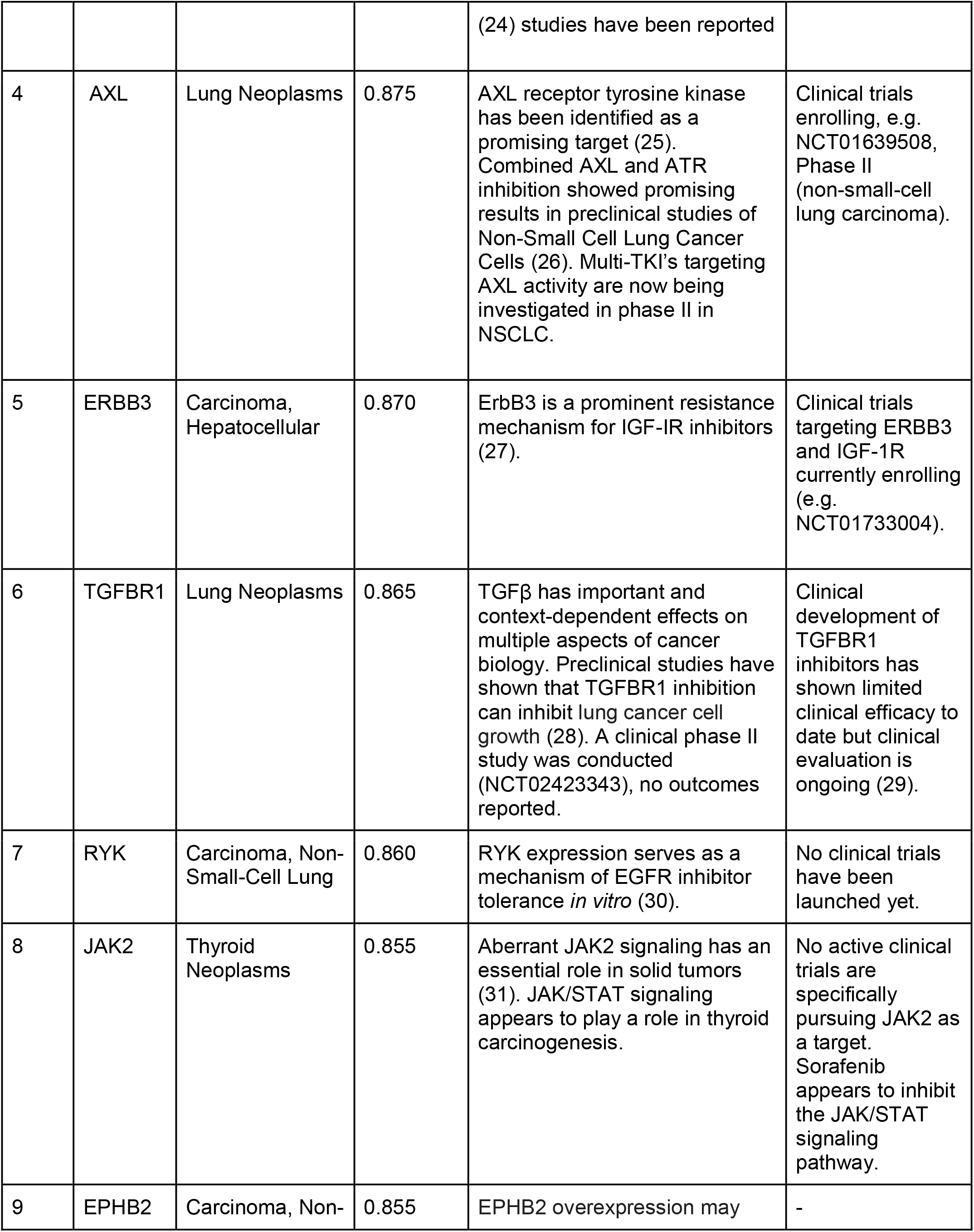
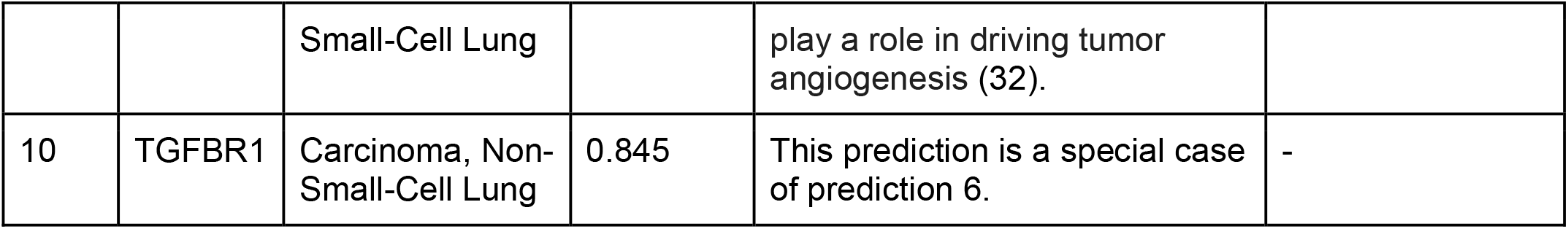
Top 10 novel PK-cancer predictions with their prediction score from the random forest classifier. These items represent predictions of PK-cancer pairs based on PubMed abstracts up to 2020 that will be found to be relevant based on phase IV PKI-cancer trials for PKIs with affinity for the PK in question.

## DISCUSSION

Phase I trials are typically performed after pre-clinical studies have suggested the potential utility of an investigational medication for a certain disease. However, less than 10% of medications entering phase I clinical testing will achieve FDA approval and reach the market (33). Therefore, there is a need for better methods for prioritizing medications and indications for clinical trials. The approach we present here does not directly prioritize PKIs and cancer indications for clinical trials, but addresses the related question of predicting whether inhibition of a given PK will be beneficial for the treatment of a given cancer.

Our work leverages word2vec (9) to generate embeddings of concepts across a large subset of abstracts in the PubMed resource as a foundation for machine learning. The key concept of word2vec goes back to the dictum of John Firth from 1957: “You shall know a word by the company it keeps” (34), meaning that context words that tend to appear near to a target word in a text corpus encode information about the word’s meaning. The embedding vectors can be regarded as compact representations of the semantic meaning of the words, which is reflected by the fact that semantically related words tend to be close to each other in the vector space, but also by the relative position of pairs of related words in the vector space. For instance, if *f* is the mapping from a large text corpus to a vector space, we often find the vectors encode similarities that capture relations between pairs of words, such as the gender relation, *f*(*woman*) - *f* (*queen*) ≈ *f* (*man*) - *f* (*king*), or the language-spoken-in relation, *f*(*Germany*) - *f*(*German*) ≈ *f*(*Italy*) - *f*(*Italian*)(35).

Our approach is motivated by the observation of the authors of word2vec that word embeddings can capture not only similarities between words (e.g., the word *France* is similar to *Italy* and other countries), but also can identify semantic relations between words (e.g., country -capital city; country - currency, adjective-adverb) (9). The basic idea of our algorithm is that these relations can relate entities of two different sets to one another, but that only some potential relations are true. For instance, the relation country - capital city is a mapping from the set of countries to the set of capital cities. The relations *France - Paris* and *Italy - Rome* are true, but the relation *France - Rome* is false.

In the biomedical sciences, there are myriad relations where we know of a limited number of true relations but are striving to identify the complete set of true relations. For instance, inhibition of PK activity has proved to be an effective anti-cancer treatment, but it is not true that inhibiting an arbitrary PK is an effective treatment for an arbitrary cancer. Only a subset of all potential pairs of PKs and cancers are true in the sense that inhibiting the PK will effectively treat a cancer. If we could accurately predict such pairs, then one could focus efforts on performing clinical trials for PKIs that inhibit the most relevant PK-cancer pairs.

Word embedding methods can represent an entire vocabulary of words into a relatively low-dimensional vector space, where semantic similarities between words are preserved in the corresponding embedded linear space (9). The embedded vectors generated by word2vec can be used as input for classification algorithms (36–38). Vector cosine similarity was taken recently to enable prediction of materials for applications years before their publication in the materials science literature by unsupervised word embedding (39). Several supervised analogy learning methods based on word embeddings have been successfully applied in a variety of natural language processing tasks (13,40,41). Our algorithm uses this approach to leverage information about cancer and kinases latent in the published literature.

Our methodology can be extended to other biomedical research questions that can be framed as a search for valid relations between concepts from two different sets. The word2vec step could be replaced by other, more advanced word embedding methods such as Bidirectional Encoder Representations from Transformers (BERT) (42) or versions of BERT such as SciBERT that are trained on scientific literature (43). In the current project, we filtered PubMed abstracts according to whether they mention cancer or PKs, but other more sophisticated relevance prediction algorithms could be employed.

## Limitations

The algorithm presented here aims to classify PKs and cancer pairs in which inhibition of the kinase can have beneficial effects in treating a certain cancer. It does not aim to predict suitability of PKIs for individual patients, which may be complicated by many factors such as the acquisition of resistance to a certain targeted treatment or genetic differences.

All phase IV studies are post-FDA approval, but not all FDA-approved drugs undergo phase IV studies. Thus, our predictions may be conservative. ClinicalTrials.gov data does not contain many studies performed outside of the US, and thus our training data may be incomplete. There is currently no standardized database with the current status of all PKIs with the results of clinical trials for the cancers they have been used to treat.

## CONCLUSION

This work presents a novel approach to predict new associations between PKs and cancers, meaning that by targeting the PKs, the corresponding cancers could be treated. We first used a word embedding algorithm to map words of the PubMed abstracts to vectors and then trained a Random Forest classifier on the embedded vectors of known pairs of PKs and cancers obtained from Clinical Trials data and Drug Central data to predict new pairs of PKs and cancers. We assessed our method using historical prediction and obtained on average AUROC above 0.8. We then applied our method on our entire corpus of PubMed abstracts and also all known pairs of PKs and cancers that were available, to predict novel pairs of PKs and cancers. From the novel predictions, we found new associations between PKs that have not been targeted yet and certain types of cancers.

## METHODS

### Text normalization and preprocessing

We developed a software package called marea (**m**area **a**damantly **r**esists **e**gregious **a**cronyms) that implements all necessary natural language processing to prepare the titles and abstracts of PubMed articles as input for word embedding algorithms. marea filters PubMed articles for relevance and applies PubTator Central (44) concept recognition to the titles and abstracts of relevant articles. After concept replacement, the final phase eliminates punctuation and stop words, and reduces the vocabulary size.

#### Filtering relevant PubMed articles

NCBI’s FTP site makes available gzipped XML files containing titles, abstracts, and metadata for all PubMed articles. marea downloads the annual baseline and daily update files, and parses them to extract the fields of interest for each article: PubMed ID, MeSH descriptors (if any), keywords (if any), and year of publication. For entries that have multiple dates with different years, the earliest one is recorded. To select articles for a particular search, the marea user provides a set of high-level MeSH descriptor ids. The MeSH descriptors defining the scope of the research described herein were D009369 (Neoplasms) and D011494 (PKs). Any article marked with at least one of these descriptors or any subcategory of these descriptors is considered relevant. An article is also judged relevant if it has a keyword that matches a label or synonym of the search descriptors or their subcategories. Some PubMed articles have neither MeSH descriptors nor keywords; some have no abstract. Any article that has no abstract is irrelevant for the search regardless of its MeSH descriptors or keywords.

#### Concept replacement

The original word2vec method operates on individual words (tokens). However, many medical concepts span multiple tokens. For instance, *non-small-cell lung carcinoma* would be treated by word2vec as three or five tokens (depending on how the hyphen is handled), but it represents a single medical concept. For this reason, recent approaches collapse multi-word concepts into a single token prior to embedding by replacing the multiword concepts with a single concept id (45). For instance, *non-small-cell lung carcinoma* can be replaced by its MeSH id D002289.

PubTator Central from the National Center for Biotechnology Information (National Library of Medicine) offers data for concept recognition in PubMed articles. Annotated categories include chemicals, diseases, genes, cell lines, SNPs, and species, as well as other categories marea does not track, such as DNAMutation and ProteinMutation. Using PubTator Central character offsets, the software replaces each phrase recognized in the title or abstract with the identifier of the corresponding concept. Diseases and chemical names are normalized to MeSH ids, genes and proteins to NCBI Gene ids, cell lines to Cellosaurus (46), SNPs to dbSNP RS ids, and species to NCBI Taxonomy ids. The one exception is the human species, NCBI taxon 9606, which we decided to skip. PubTator Central annotations would have substituted 9606 for *man, woman, boy, girl, father, mother, patient*, and similar words. We chose to preserve the distinctions of gender and age expressed in terms for humans, as these factors are certainly significant in the medical context.

#### Text preprocessing

After concept replacement, marea cleans up the text of PubMed titles and abstracts to make it more suitable for word embedding. The tokenizer deletes all punctuation symbols, including hyphens and underscores within words: the parts of a compound word become separate tokens. marea removes stop words, whether lowercase or capitalized. Uppercase acronyms of length ≥ 2, even those that coincide with stop words, are not changed. For example, the acronym *ALL* (acute lymphocytic leukemia) is retained while *all* and *All* are eliminated. We started with the stop word list for English in the Natural Language Toolkit (nltk version 3.5) Python library (47) and added some new stop words. Any letter of the alphabet that occurs as a single-character token is a stop word. To further reduce the size of the vocabulary, tokens that remain after stop word removal are lemmatized with the WordNet (48) lemmatizer from nltk. The lemmatizer reduces words to their base form, for example plural nouns are simplified to the singular. (Unlike stemming, lemmatizing a word always returns a complete word, not a truncated word stem.) The last step of text preprocessing converts everything to lowercase, to avoid near-duplicate embeddings for upper-, lower-, and mixed-case forms of the same word.

### Word embedding

The word embedding method based on the word2vec algorithm is performed on the preprocessed corpus to embed words to vectors. We used the EMBeddInG GENerator (embiggen), a Python 3 software library developed by our group for word embedding based on word2vec and node embedding based on the node2vec algorithm (49). In the current project, the skip-gram model was used for word2vec with the parameters window size = 5, minimum count (minimum word frequency) = 5, batch size = 128, negative samples = 20 and dimension = 100. Word embedding on the total corpus resulted in embeddings of 293,274 words each with dimension 100.

### PKIs and their PK targets

We used the online drug compendium DrugCentral (50) to explore all the kinase activities. DrugCentral keeps track of all known experimental activities for approved drugs across all major protein target families (including kinases). Hence, we extracted the kinase activities from DrugCentral and matched them to the drugs that are PKIs only. The result of this operation is a list of PKI-PK pairs (PKI2PK), each of which is mapped to an experimental value of affinity (e.g. Ki, IC_50_, etc) in micromolar units and appropriately referenced (when possible) with a PubMed ID (PMID). Moreover, we kept only the PKI2PK pairs having an activity value below 0.03 µM, which is the threshold under which drugs are more likely to act on kinases (51). The last filter that we applied to extract PKI2PK pairs was the number of PKs that are inhibited by a PKI to treat a cancer. For our analysis, we chose PKIs that have an affinity value below 0.03 µM and inhibit at most 5 PKs. If a PKI inhibited more than 5 PKIs at this threshold, we chose the top five PKs. Filtering the DrugCentral data by applying the affinity threshold 0.03 µM and the count of PKs targeted by a PKI to 5 resulted in a list of 226 pairs of PKs and PKIs which is given in **Supplementary Material File 1**. The MeSH sub-hierarchy that descends from D009369 (Neoplasms) includes experimental (rodent) cancer models, veterinary cancer types, and also non-specific categories. These were removed prior to further analysis (both **Table 1** and **Supplementary Material File 3**).

### Phase I-Phase IV clinical trials of PKIs for cancer therapy

Clinical trials are typically performed in four standardized phases. A phase I trial is designed to test the safety and pharmacology of a drug. Phase II trials are therapeutic exploratory trials that are conducted in a small number of volunteers who have the disease of interest, and to answer questions required to prepare a phase III trial including optimal doses, dose frequencies, administration routes, and endpoints. Phase III trials strive to demonstrate or confirm efficacy, often by comparing the intervention of interest with either a standard therapy or a placebo. Additionally, the incidence of common adverse reactions is characterized. Phase IV trials are performed subsequent to initial FDA approval with the goal of identifying less common adverse reactions and in some cases of evaluating a drug in populations different from the original study population (52).

We downloaded the Clinical Trials data from the ClinicalTrials.gov server and also obtained the list of all MeSH terms that descend from “Neoplasms”. This list contains 698 neoplasms and their MeSH ids. Using the Clinical Trials data and list of neoplasms and their MeSH ids, we created a list of neoplasms and PKIs that were used to treat the cancers along with the clinical trial phase, start date, completion date of the clinical trials study, MeSH id for each neoplasm and NCT id for each clinical trial study. This dataset is presented in the **Supplementary Material File 2**.

### Historical validation

In order to estimate the performance of our approach, we trained our model on historical snapshots of PubMed and tested the predictive accuracy with Clinical Trials data from subsequent years. For each experiment, we fixed the target year to a specific year and used PubMed abstracts published up to and including this year for word embedding. We constructed the positive and negative training sets described below but limited the Clinical Trials data to entries that were initially published not later than the target year.

To create the positive training set, we chose all pairs of PKs and cancers where the PKIs were approved to treat the cancers in the phase IV of the Clinical Trials data up to a target year. To create the negative training set, we randomly generated pairs of PKs and cancers where there was no evidence of treating the cancers by inhibiting the PK in the Clinical Trials data up to the target year. The negative test set was defined based on randomly chosen PKs and cancers for which no trial of any phase was present in the Clinical Trials data up to the target year. Also, there was no PK-cancer pair in common between the negative training set and negative test set. In our implementations, we fixed the size of the negative training/test set to be ten times of the size of the positive training set. However, the size of the generated negative training/tes set might be smaller, if the vectors corresponding to the randomly chosen PKs or cancers are not found in the embeddings and it would not be possible to generate any other PK-cancer pair.

In our first experiments, the positive set was defined on the basis of phase I, II, III, and IV studies, i.e. it contained pairs of PKs and cancers where the PKIs were approved to treat the cancers in at least phase I of the Clinical Trials data after the target year. However, we made sure that there was no evidence of inhibiting the PK in the Clinical Trials data in any phase until the target year. In our second experiments, we defined the positive set on the basis of only phase IV studies. Again, we made sure that there was no evidence of inhibiting the PK in the Clinical Trials data in any phase in the target year and before that.

### Random forest learning

The next step after generating positive/negative training/test sets which contain lists of PKs-cancer pairs is to find the embeddings of PKs and cancers and prepare the datasets for the prediction task. For a given pair of a PK and a cancer, we subtracted the vector corresponding to the cancer from the vector corresponding to the PK. The difference vectors from the positive training and test sets were labeled with 1 and the difference vectors from the negative training and tests were labeled with 0.

Random forest learning was performed in Python 3.7, using scikit learn 0.24.1. A randomized search was performed on different parameters including number of estimators, maximum features, maximum depth, minimum samples split, minimum samples leaf and bootstrap and also 10-fold cross validation. The best model was selected for the prediction task.

### Performance assessment

The results of predictions are demonstrated as the area under curve (AUC) scores of the Receiver Operating Characteristic (ROC) curves (AUROC). AUROC is a measure of the ability of the classifier to distinguish between the two classes (PK-cancer pairs and non PK-cancer pairs).

### Source code

Several code repositories were developed for this project. **marea** performs concept replacement and preprocessing of PubMed abstracts and is available at https://github.com/TheJacksonLaboratory/marea under the BSD 3 license. Yet another clinical trials parser (YACTP) retrieves and processes information from ClinicalTrials.gov and is available at https://github.com/monarch-initiative/yactp under the GNU General Public License v3.0. Kinase Cancer Embedding Tool (KCET) is available at https://github.com/TheJacksonLaboratory/KCET and contains scripts and Jupyter notebooks used to perform word embedding and to leverage the embeddings for random forest classification. The analysis described in this manuscript corresponds to release v0.2.0. The embedding software, embiggen, performs word embedding and is available at https://github.com/monarch-initiative/embiggen as well as via PyPi at https://pypi.org/project/embiggen/.

## ACKNOWLEDGMENT

This work was supported by NIH grant U01-CA239108-02 (Illuminating the Druggable Genome by Knowledge Graphs). The authors would like to thank Karen Davis for assistance in creating the figures.

