## Supplemental Material for "Supervised learning with word embeddings derived from PubMed captures latent knowledge about protein kinases and cancer"

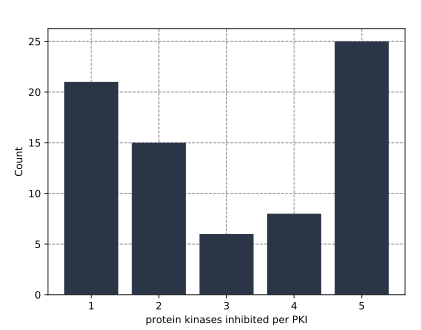

**A)**

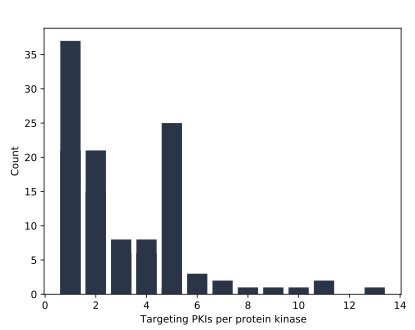

**B)**

**Figure S1**. **A)** The histogram of the number of PKs that are inhibited by a given PKI,

**B)** The histogram of the number of PKIs that inhibit a given PK.

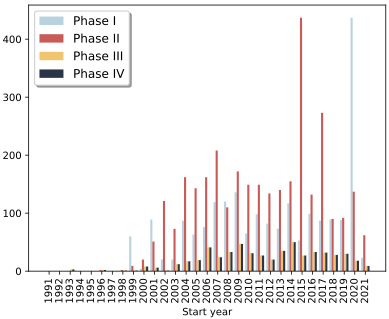

**A)**

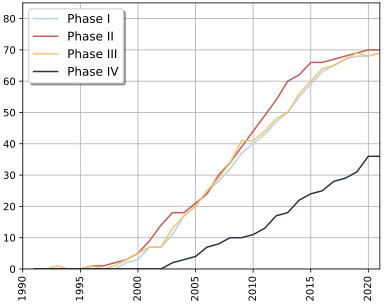

**B)**

**Figure S2**. **A)** Histogram of phases I, II, III and IV of the clinical trials data from 1991 to 2021.

**B)** Number of PKIs being studied per year and per phase from1991 to 2021.

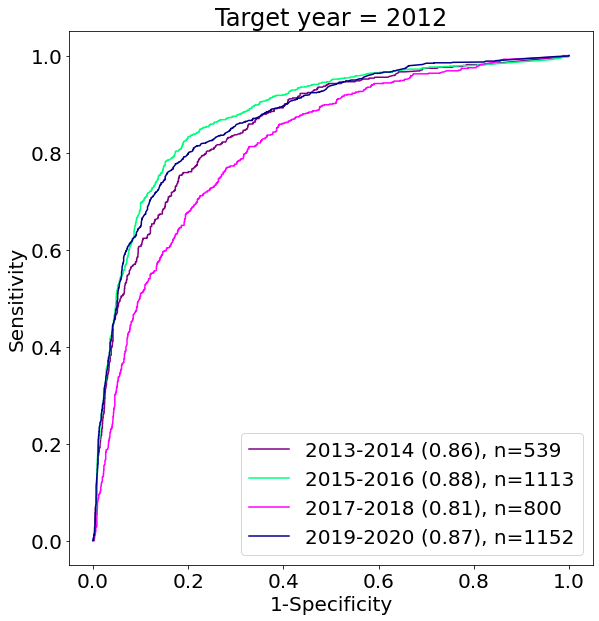

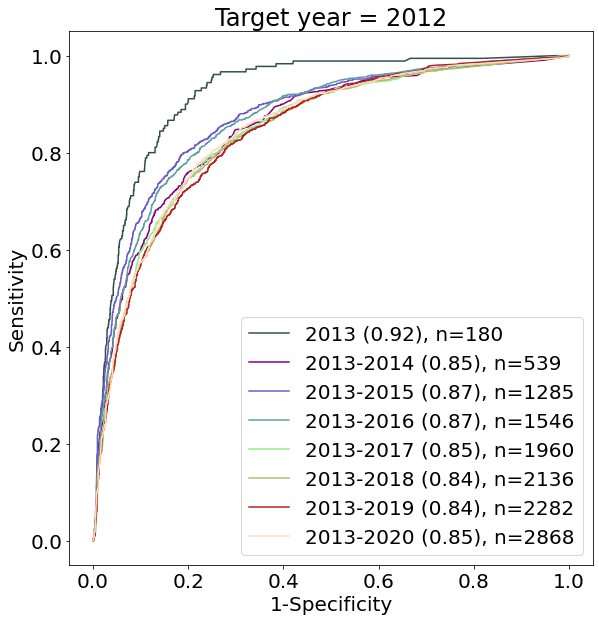

A) **B)**

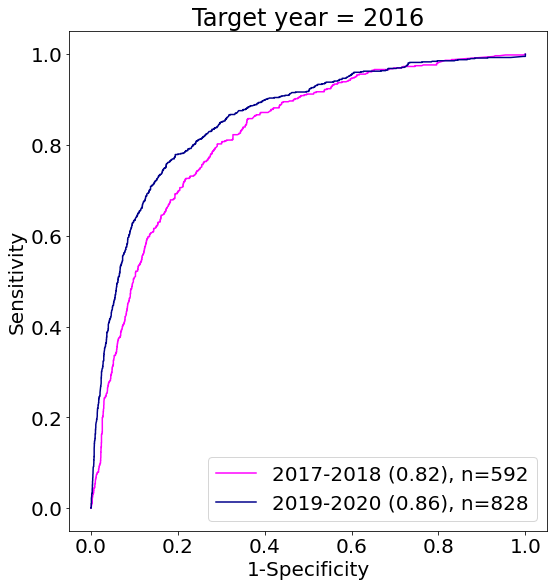

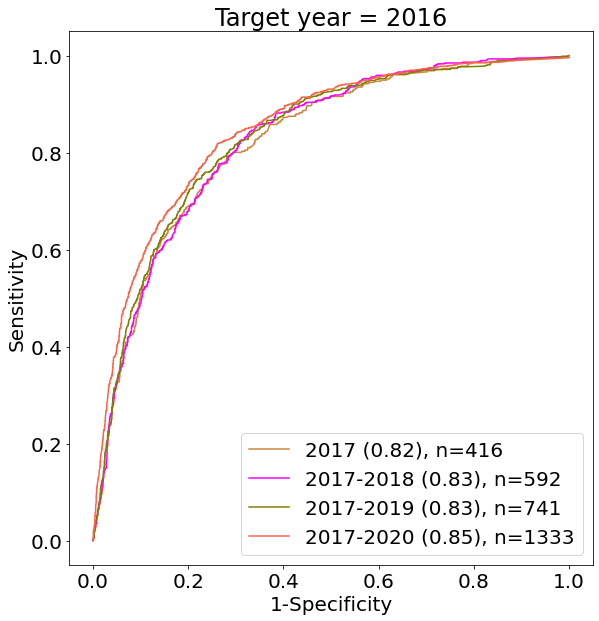

**C)	D)**

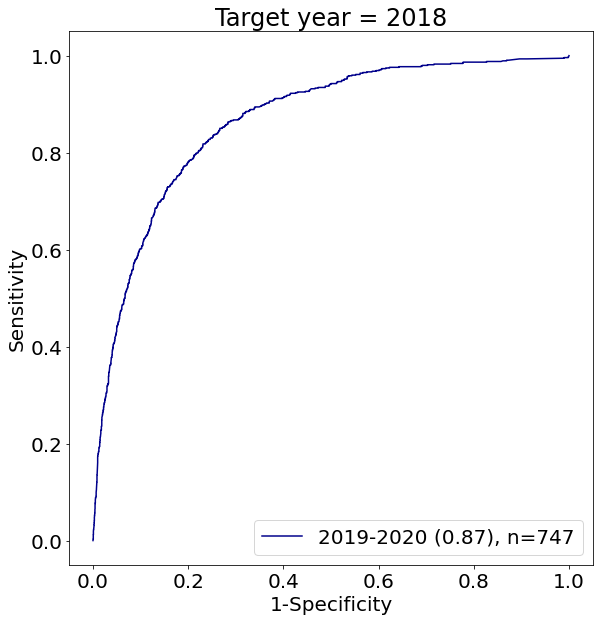

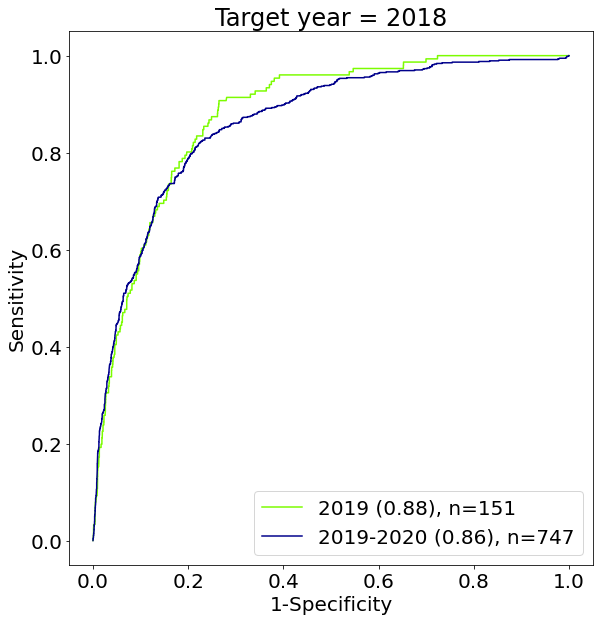

**E)	F)**

**Figure S3**. **ROC analysis of predicted PK-cancer pairs (all clinical trial phases).**

**A)** Predictions based on abstracts published up to 2012, grouped according to two year periods following 2012. **B)** Predictions based on abstracts published up to 2012, grouped according to increasing periods of time in the future. **C)** Predictions based on abstracts published up to 2016, otherwise analogous to panel A. **D)** Predictions based on abstracts published up to 2016, otherwise analogous to panel B. **E)** Predictions based on abstracts published up to 2018, otherwise analogous to panel A. **F)** Predictions based on abstracts published up to 2018, otherwise analogous to panel B.

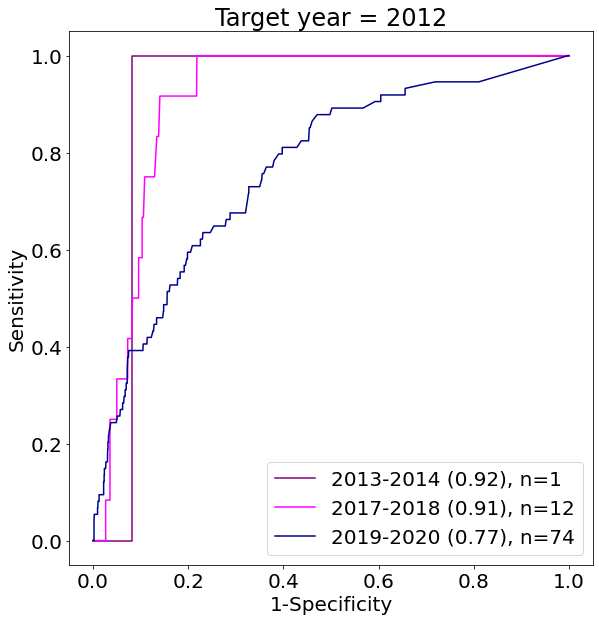

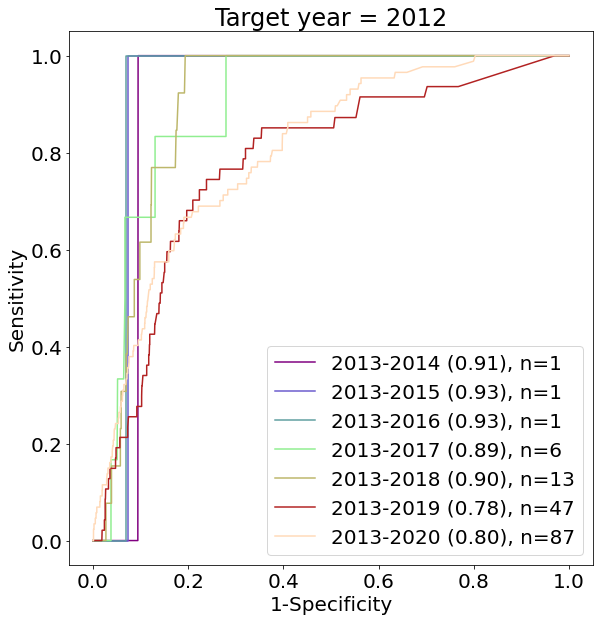

A) **B)**

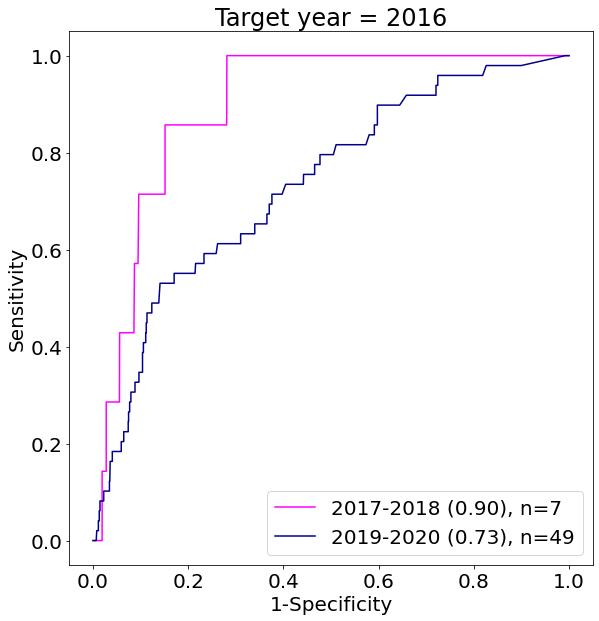

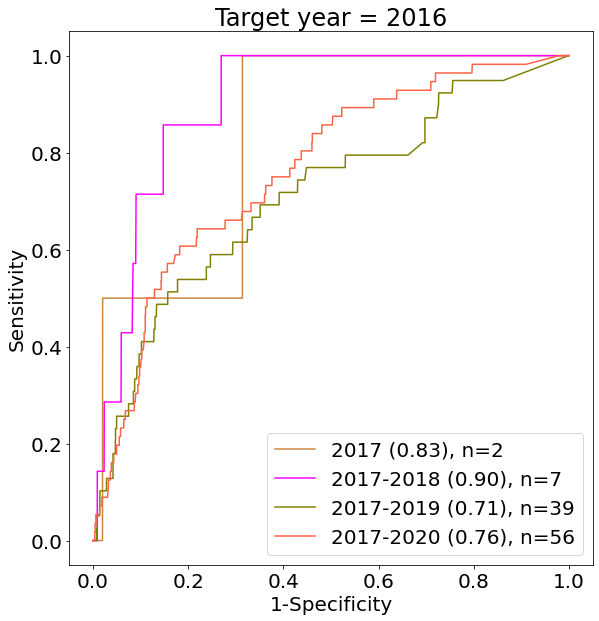

**C)	D)**

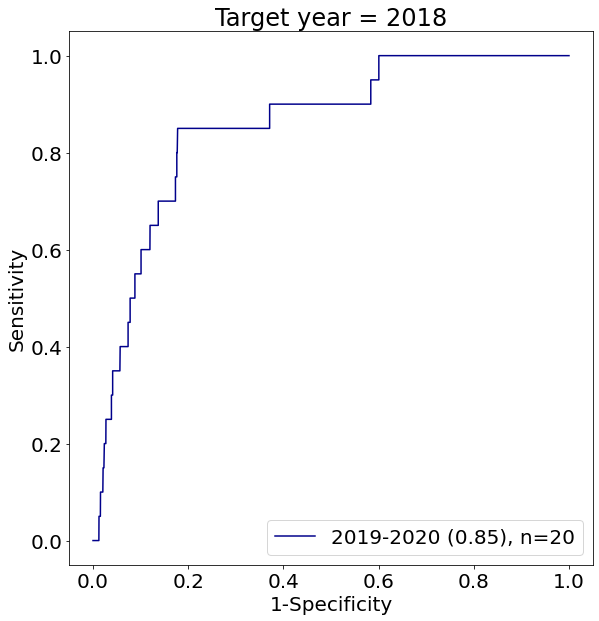

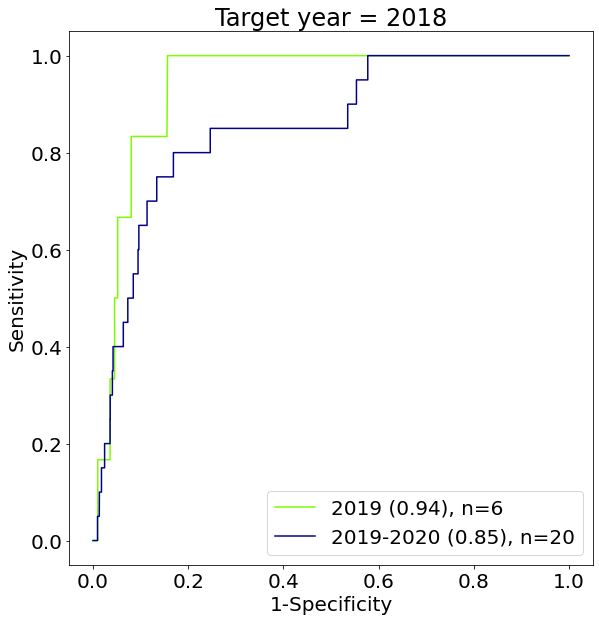

**E)	F)**

**Figure S4**. **ROC analysis of predicted PK-cancer pairs (phase IV clinical trials).**

**A)** Predictions based on abstracts published up to 2012, grouped according to two year periods following 2012. **B)** Predictions based on abstracts published up to 2012, grouped according to increasing periods of time in the future. **C)** Predictions based on abstracts published up to 2016, otherwise analogous to panel A. **D)** Predictions based on abstracts published up to 2016, otherwise analogous to panel B. **E)** Predictions based on abstracts published up to 2018, otherwise analogous to panel A. **F)** Predictions based on abstracts published up to 2018, otherwise analogous to panel B.
